# Distinct contributions by frontal and parietal cortices support working memory

**DOI:** 10.1101/104430

**Authors:** Wayne E. Mackey, Clayton E. Curtis

## Abstract

Although subregions of frontal and parietal cortex both contribute and coordinate to support working memory (WM) functions, their distinct contributions remain elusive. Here, we demonstrate that perturbations to topographically organized human frontal and parietal cortex during WM maintenance cause distinct but systematic distortions in WM. The nature of these distortions supports theories positing that parietal cortex mainly codes for retrospective sensory information, while frontal cortex codes for prospective action.

**Acknowledgements:** The US National Institutes of Health (R01 EY016407 to C.E.C.) and the National Science Foundation (Graduate Research Fellowship Program to W.E.M.) supported the research.

**Author Contributions:** W.E.M and C.E.C. designed the experiment, collected and analyzed the data, and wrote the manuscript.

**Conflict of interest:** The authors declare no competing financial interests.

## Main text

Working memory (WM) bridges perception and action over brief periods of time ^1^ and acts as a critical building block for high-level cognitions ^2–4^. While a large network of brain areas support WM ^5^, past research demonstrates that within portions of frontal and parietal cortex, specifically, the activities of neurons persist during WM retention intervals ^6–8^. Moreover, chronic lesions to both of these areas impair WM ability ^9–12^. The patterns of neural activity during WM and the consequences of damage are so strikingly similar, it remains unknown what distinct contributions, if any, the frontal and parietal cortices provide to support WM.

We addressed this key limitation by testing the hypothesis that the nature of the maintained information differs in each area. Based on theories of sensorimotor dynamics ^1, 13^, we predict that the parietal cortex largely maintains representations of past sensory information, while the frontal cortex largely maintains representations of future plans. We test this hypothesis by combining computational neuroimaging and transcranial magnetic stimulation (TMS) to transiently disrupt activity in topographically defined subregions of frontal and parietal cortices while subjects actively maintained information in WM.

We used functional magnetic resonance imaging (fMRI) and nonlinear population receptive field (pRF) mapping (**Fig. 1a,b,**) ^14, 15^ to identify potential stimulation sites in individual subjects, including four visual field maps in parietal cortex (IPS0, IPS1, IPS2, and IPS3) and two visual field maps in frontal cortex (SPCS, IPCS) (**Fig. 1c,d**). We targeted the superior precentral sulcus (sPCS) in frontal cortex and the third intraparietal sulcus area (IPS2) in parietal cortex because those topographic maps are the putative human homologues of the monkey frontal eye field (FEF) and lateral intraparietal area (LIP), respectively. In both species, these areas exhibit delay-period activity during WM tasks ^6, 8, 16^, and lesions to these areas in both species cause impairments in WM ^10–12, 17, 18^. Additionally, we targeted the dorsolateral prefrontal cortex (PFC), an area of frontal cortex also shown to be critical for WM in nonhuman primates ^9^, but whose role in human WM is controversial ^12^. Consistent with previous studies, we found no coherent topographic map of visual space in dorsolateral PFC ^15, 19, 20^, and therefore used anatomical landmarks for criteria to identify dorsolateral PFC in individual subjects ^21^.

**Figure 1.**
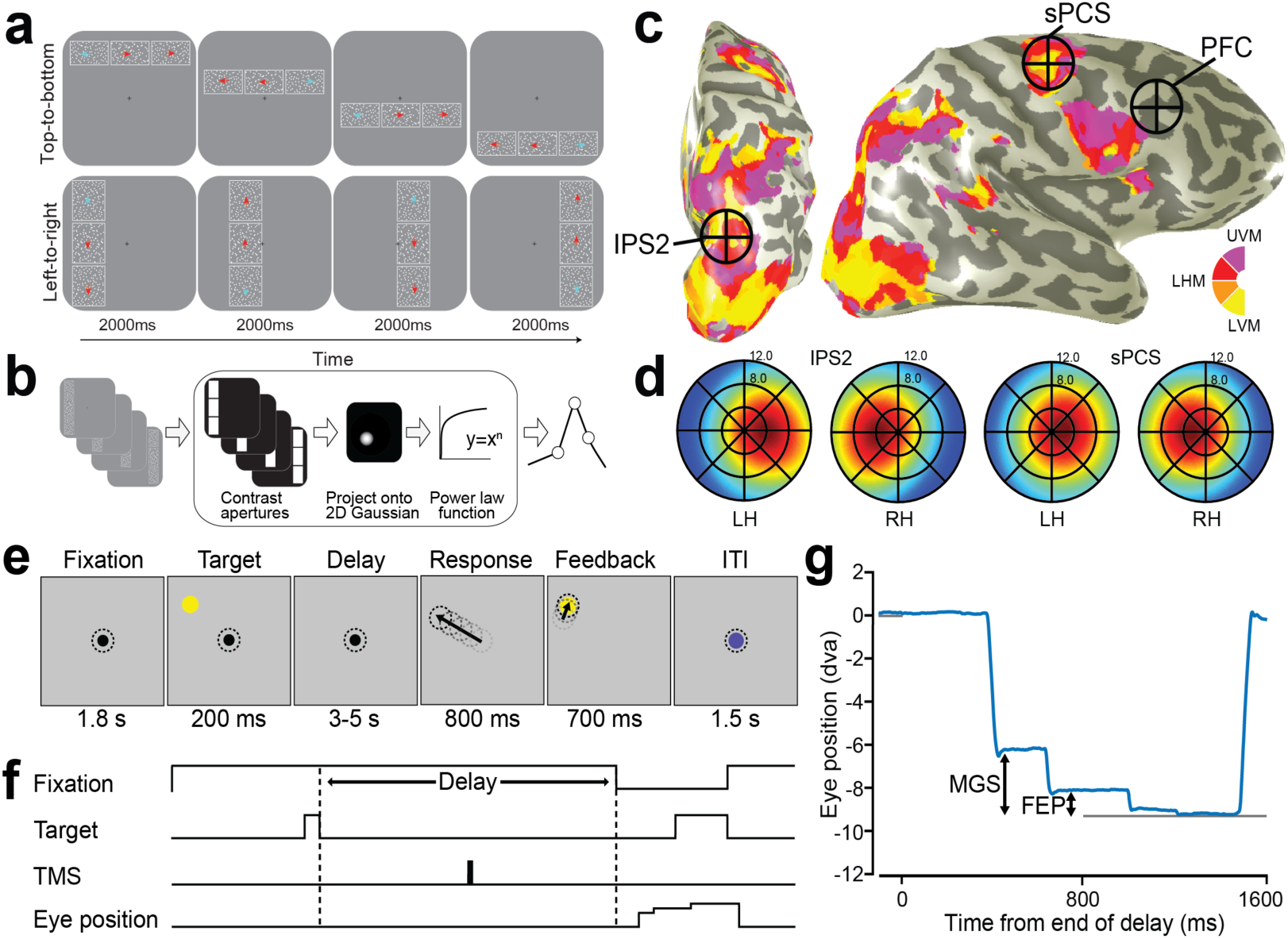
Experimental procedures. (***a***) Discrimination task used for topographic mapping. Subjects fixated centrally while covertly attending to a bar consisting of three apertures of moving dots that swept across the screen. Subjects pressed a button to indicate which flanking aperture (left or right; above or below) contained dots moving in the same direction as the center sample aperture. Staircases adjusted dot motion coherence in the flanking apertures to constantly tax attention. White outlines around each aperture were not visible to the subject, but are shown here for clarity. (***b***) Nonlinear population receptive field (pRF) modeling schematic. Stimulus sequences from the discrimination task were converted into 2D binary contrast apertures and projected onto an isotropic 2D Gaussian that represented a predicted pRF. Static nonlinearity was applied to account for compressive spatial summation. (***c***) Stimulation sites projected onto the inflated cortical surface of a sample subject. The color indicates the best-fit phase angle by the pRF model. sPCS and IPS2 stimulation sites were localized by our mapping procedure, and dorsolateral PFC stimulation targets were defined by individual subject anatomical landmarks. (***d***) Visual field map coverage in IPS2 and sPCS. While visual fields maps in both regions primarily represent the contralateral hemifield, they also represent portions of the ipsilateral hemifield. (***e***) Memory-guided saccade task. While fixating, subjects maintained the position of a brief visual target over a memory delay, and then made a saccade to the remembered target. The correct target location was again presented for feedback. Dotted circles depict gaze, but were not visible to subjects. (***f***) Temporal structure of the experimental task and application of TMS. In order to disrupt the maintenance of information, we applied a short burst of theta burst stimulation (TBS) to IPS2, sPCS, or dorsolateral PFC during the middle of the delay period on each trial. (***g***) An example horizontal eye position trace shows two distinct types of memory error relative to the true target location: the endpoint of an initial memory-guided saccade (MGS), and the final eye position (FEP) following quick corrective saccades (in this example only one) prior to the re-presentation of the target feedback (gray line).

We measured the degree to which disrupting neural activity with TMS in each target area affected WM performance, which was assayed by the accuracy of memory-guided saccades (**Fig. 1e**). Subjects generated saccades following a memory retention interval to the location of a target briefly flashed before the delay. During TMS sessions, but not control sessions, we applied short trains of theta-burst TMS in the middle of the delay period on each trial (**Fig. 1f**). Subjects typically make an initial ballistic eye movement called a memory-guided saccade (MGS) towards the remembered target, followed by small and sometimes multiple corrective eye movements that usually bring the gaze closer in alignment with the remembered target (**Fig. 1g**). Therefore, we measured the accuracies of both the initial MGS and the final eye position (FEP) on each trial. Based on previous work, we know that MGS and FEP are sensitive to different forms of oculomotor adaptation ^22^, have different developmental trajectories ^23^, and different patterns of dysfunction in schizophrenia ^24^. More importantly, they may reflect different components of WM ^11, 12^. The accuracy of the initial MGS may index the quality of the prospective movement, in this case the saccade plan. The FEP, on the other hand, may be an indicator of the fidelity of retrospective sensory information. These two WM codes differ in the nature of what is maintained in memory, a future action plan or a past sensory event.

TMS disruption in frontal and parietal cortex caused distinct WM impairments (**Fig. 2**). Although disruption to sPCS caused increased errors in MGS to the contralateral visual field (p=0.005; TMS worsened MGS in every subject), and to a lesser extent to the ipsilateral visual field (p=0.02), it had no effect on the accuracy of FEP (contralateral p=0.54, ipsilateral p=0.74). Disruption to IPS2 caused increased errors in FEP in the contralateral visual field (p=0.0005; TMS worsened FEP in every subject), but not ipsilateral visual field (p=0.24). Similar to sPCS, IPS2 disruption also increased MGS errors to the contralateral visual field (contralateral p=0.04; ipsilateral p=0.28). The pattern of results is consistent with the assertion that frontal cortex maintains a code of prospective action, while parietal cortex maintains a code of retrospective perception. However, this is not true of the entire frontal cortex. Disruption of dorsolateral PFC, just a few centimeters away from topographically organized sPCS, caused no observable impairments in MGS (contralateral p=0.32, ipsilateral p=0.74) or FEP (contralateral p=0.37, ipsilateral p=0.67), consistent with the effects recently reported from chronic dorsolateral PFC lesions ^12^.

**Figure 2.**
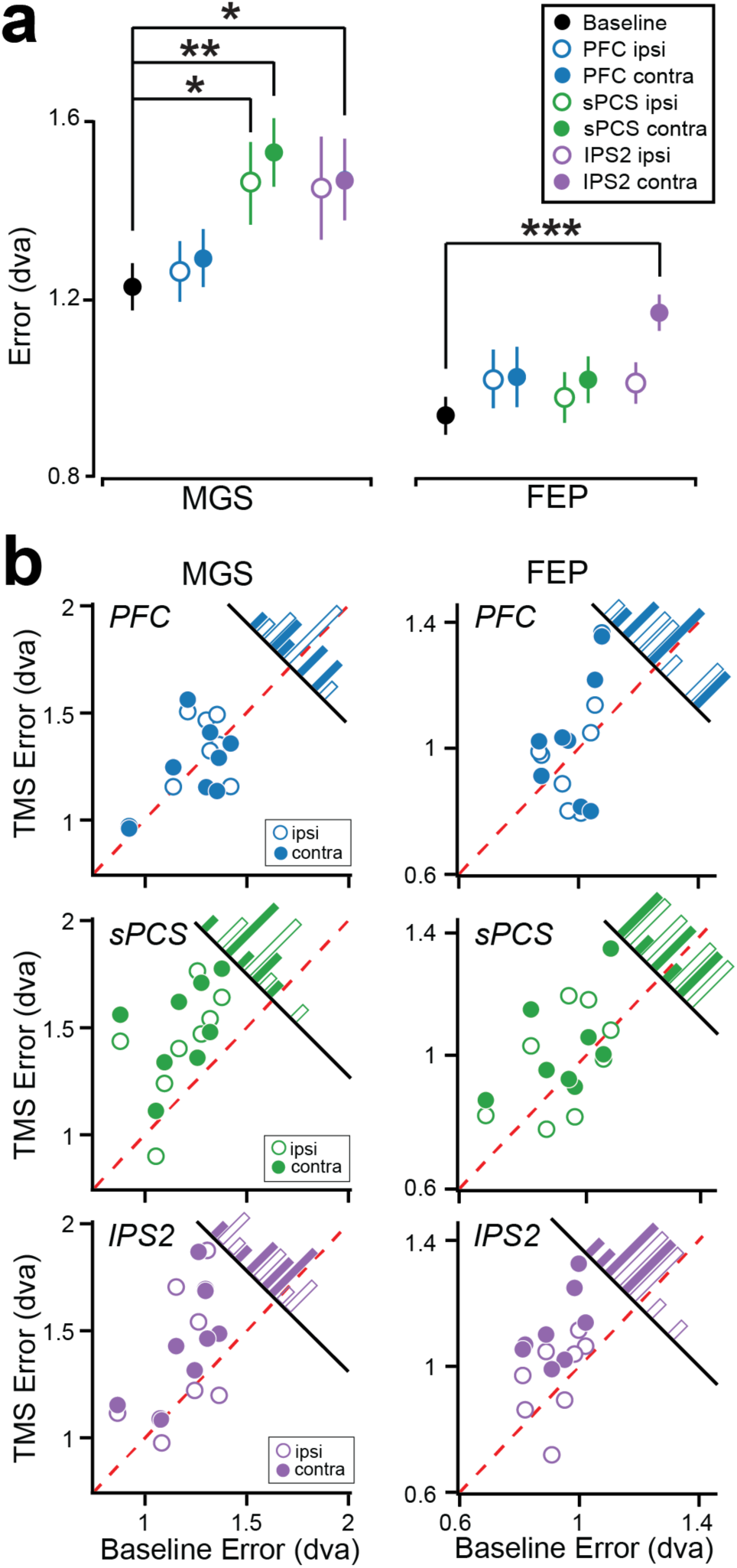
Group average and individual working memory performance. (***a***) Compared to a baseline no TMS condition, sPCS TMS caused an increase in mean MGS errors especially in the visual field contralateral to the hemisphere of TMS, but not FEP errors. IPS2 TMS impaired mean FEP errors in the contralateral visual field. Dorsolateral PFC TMS caused no observable impairments. ✝ p <=0.08, * p <=0.05, ** p <=.005, *** p <=0.0005. Error bars represent standard error of the mean. (***b***) Distribution of effects of TMS on working memory performance. TMS induced error (y-axis) is plotted against baseline error (x-axis) for both MGS and FEP, following PFC, sPCS, and IPS2 TMS. Histograms show the distributions of TMS induced error. Note that individual performance is remarkably consistent across subjects, and parallels the group-level results. For instance, sPCS TMS caused increased MGS error compared to baseline for every subject (all dots are to the left of the red dashed identity line). Additionally, IPS2 TMS caused increased FEP errors in every subject.

These results in the context of previous work have three important implications. First, although lesions to the macaque dorsolateral PFC severely impair the accuracy of memory-guided saccades ^9^, we recently demonstrated that surgical resections of human dorsolateral PFC had no impact on memory-guided saccade accuracy ^12^. Our current results also indicate that TMS disruption of human PFC has no impact on WM performance, and thus rule out the possibility that cortical reorganization compensated for our previous patient results. Moreover, they point to a fundamental difference between the two species in terms of the necessities of human and monkey dorsolateral PFC for WM. Second, we found that TMS disruptions targeting retinotopic portions of the precentral sulcus and intraparietal sulcus caused systematic impairments in every subject. Unlike PFC, these human results perfectly mirror impaired WM following perturbation or damage to macaque FEF and LIP ^9^, ^10^. More importantly, they strongly suggest that delay period activity within these frontoparietal topographic maps that we presumably disrupted with TMS maintains spatial representations used for memory-guided responses ^20^. Third, TMS disruption of precentral and intraparietal cortex caused distinct types of WM errors suggesting distinct roles in WM. Topographic patterns of neural activity maintain WM representations of space ^20^. Perturbation of sPCS maps distorted these representations causing increased errors in memory-guided saccade programming. Presumably, downstream oculomotor areas (e.g., superior colliculus) would plan the metrics of the forthcoming saccade by the read-out of a corrupted map. However, unperturbed topographic maps in parietal cortex could then be read-out to correct for the spatial mismatch between its stored location and the erroneous memory-guided saccade. TMS to the IPS2 topographic map distorted a more general representation of the past target’s location, affecting both the accuracy of the memory-guided saccade and any subsequent corrections. Thus, WM delay period activity in the frontal and parietal cortex may differentially code for prospective action plans and retrospective sensory information, respectively. Although human neuroimaging support this assertion ^25, 26^, future work should focus on addressing to what extent these distinctions are absolute or relative and what extent these codes depend on one another.

## Online Methods

### Subjects

Nine neurologically healthy individuals (2 female) with normal or corrected-to-normal vision took part in the study. All subjects gave written informed consent before participating and were compensated monetarily. All procedures were approved by the human subjects Institutional Review Board at New York University. All nine subjects completed a baseline session of the MGS task without TMS. Seven subjects completed a session of each TMS condition (PFC, PCS, IPS), while one subject only completed a PCS TMS session, and another completed PFC and IPS TMS sessions. This resulted in eight subjects in each of the three TMS conditions.

### MRI Acquisition

MRI data were collected using a 3T head-only scanner (Allegra; Siemens) at the Center for Brain Imaging at New York University. Images were acquired using a custom four-channel phased array receive coil (NOVA Medical) placed over lateral frontal and parietal cortices. Volumes were acquired using a T2*-sensitive echo planar imaging (EPI) pulse sequence [repetition time (TR), 2000 ms; echo time 30 ms; flip angle, 75°; 31 slices; 2.5 mm × 2.5 mm × 2.5 mm voxels]. Low resolution T1-weighted anatomical images were collected at the beginning of each scanning session using the same slice prescriptions as the functional data. High-resolution T1-weighted scans (1 mm × 1 mm × 1 mm voxels) were collected for registration, segmentation, and display.

### Topographic mapping procedures

Observers performed a difficult discrimination task that required covertly attending to stimuli within bars of different widths that randomly swept across the visual field in different directions. The bars subtended 1 degree, 2 degrees, or 3 degrees of visual angle. Each bar was split into thirds. For example, a bar that swept from right to left was split into a top patch, center patch, and bottom patch. A bar that swept from top to bottom was split into a left patch, center patch, and right patch. We asked subjects to select which patch of moving dots matched the direction of motion in the center patch. In order to keep the discrimination task difficult, we used a two up one down staircase on the coherence value for the moving dots. Stimuli were generated in MATLAB with the MGL toolbox and displayed on a screen in the bore of the magnet. Subjects viewed the display via a mirror mounted on the RF coil. Behavioral responses were recorded using a button box.

### MRI Preprocessing

T1-weighted anatomical scans were automatically segmented using Freesurfer. All fMRI analysis was performed using mrVista (http://vistalab.stanford.edu/software). The first three volumes of each functional run were removed to allow magnetization to reach a steady-state. Subsequent volumes were slice-time and motion corrected. Data was then aligned to each individual subject’s T1-weighted anatomical image. Functional scans for each individual experimental bar size (1, 2, and 3 degrees) were averaged together separately. Cortical surfaces were reconstructed at the gray/white matter border and projected as a smoothed 3D mesh or flattened cortical representation.

### PRF Analysis

We modeled response amplitudes for each voxel using a modified version of the pRF model described by Dumoulin & Wandell (2008) that incorporates a static power-law nonlinearity to account for non-linear compressive spatial summation ^27^. This model allows us to estimate an individual voxel’s receptive field center and size. Typically, the pRF model consists of an isotropic gaussian with four parameters: position ( *x, y* ), size (σ), and amplitude (*β*). The CSS model we employed adds an additional parameter, an exponent (*n*). This model is expressed formally as:

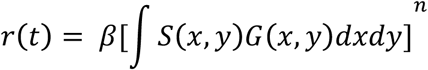

where *r(t)* is the voxel’s predicted response, *S* is the binary stimulus image, and *G* is an isotropic gaussian expressed as:

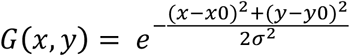

We have previously shown that the CSS pRF model is more accurate than the conventional linear pRF model, especially in parietal and frontal cortex ^15^.

### Oculomotor procedures

Monocular eye-movement data were collected at 1000 Hz using an SR Research EyeLink 1000 eye-tracker. Subjects sat in a darkened room while their head was stabilized using a chin rest to eliminate head movement and help them remain comfortable during the task. Nine point calibrations were performed at the beginning of each session as well as between runs whenever necessary. Experimental stimuli were displayed against a gray background and programmed in MATLAB (The MathWorks) using the MGL toolbox.

### Transcranial magnetic stimulation procedures

TMS was administered using a Magstim Rapid 2 Magnetic Stimulator (Magstim) with a figure-eight coil (70-mm diameter double circle). We stimulated left sPCS and left IPS2 as defined by our topographic mapping experiment. We stimulated right dorsolateral PFC, defined as the posterior third of the intermediate frontal sulcus ^21^. We chose to stimulate left sPCS and IPS2 and right dorsolateral PFC in order to strengthen the test of our hypothesis. As it has long been suggested the right hemisphere plays a privileged role in spatial processing compared to the left hemisphere, we wanted to rule out the chance that this dominance could interfere with our hypothesis. Stimulation was applied using an online theta burst protocol consisting of 1 train of 3 pulses of 50hz ^28^. Stimulation intensity was set at 53% maximum stimulator output for each subject.

### Experimental procedure

Subjects began by fixating a preparation stimulus (black cross over white dots) at the center of the screen. A target (yellow, 0.5° diameter dot) then briefly flashed (200 ms) at a random location of 10 degrees eccentricity from the central fixation point. No targets were presented near the cardinal axes to prevent verbalizing of locations (e.g., up, down). We instructed subjects to remember the location of the dot for a variable delay period (3, 3.5, 4, 4.5, or 5 s). At the end of the delay period, a sound coupled with the disappearance of the fixation point instructed the subject to shift their gaze to the spatial location of the target they were holding in memory. After 800 ms, the target was represented on the screen (as a green dot for 700 ms) to provide feedback to the subject. They were trained to look at the target if their gaze was incorrect (i.e., displaced from the target). Afterward, an intertrial interval (blue square at central fixation, 1.5 s) was displayed to notify subjects that the current trial had ended and a new one was about to begin. Each run consisted of a total of 30 trials. All subjects completed a total of 10 runs and were encouraged to take breaks between runs as they felt necessary to remain both comfortable and alert.

### Analysis

Eye-movement data were transformed into degrees of visual angle using a third-order polynomial algorithm that fit eye positions to known spatial locations and then scored offline with an in-house MATLAB function-graphing toolbox (iEye, https://github.com/wemackey/iEye). Saccades were defined as when eye velocity exceeded 30 degrees/s or when velocity failed to reach 30 degrees/s, confirmed by visual inspection. Error was defined as distance between the response location and the target location, expressed in degrees of visual angle. Saccadic response time represents the amount of time, in milliseconds, between the onset of the response cue and saccade onset. Trials where saccadic response time exceeded 900 ms or were shorter than 100 ms were discarded from analysis. Trials where subjects prematurely broke fixation were also discarded.

We grouped results by the visual field the target location appeared in relative to the hemisphere of the stimulation site (contralateral or ipsilateral to TMS coil). Since performance in the left and right visual fields did not differ (Wilcoxon Rank-Sum) in the no TMS condition, we collapsed across those trials to estimate baseline performance. This resulted in a total of seven performance groups: no TMS, PFC-TMS contralateral, PFC-TMS ipsilateral, sPCS-TMS contralateral, sPCS-TMS ipsilateral, IPS2-TMS contralateral, and IPS2-TMS ipsilateral. Additionally, we found that performance did not differ between the narrow range of delays used (Wilcoxon Rank-Sum) and therefore collapsed our analyses across delays. We performed statistical analysis of performance results across groups (Kruskal-Wallis ANOVA) for all metrics. When results were statistically significant, we compared ipsilateral and contralateral TMS performance with baseline performance (Wilcoxon Rank-Sum).

